# Ecological insights into soil health according to the life-history traits and environment-wide associations of bacteria in agricultural soils

**DOI:** 10.1101/2022.02.03.479020

**Authors:** Roland C. Wilhelm, Joseph P. Amsili, Kirsten S.M. Kurtz, Harold M. van Es, Daniel H. Buckley

**Author notes:** Corresponding Author: Dr. Daniel Buckley, Current address: School of Integrative Plant Science, 306 Tower Road, Cornell University, Ithaca, NY 14853.

## Abstract

Soil health assessment may be enhanced by monitoring changes in bacterial populations that are indicators of various biological, physical, and chemical properties of soil. However, the lack of ecological information for many abundant bacteria in agricultural soils limits our understanding of indicator responses and, thus, their utility for guiding management. We identified bacterial indicators of twelve conventional measures of soil health, and tillage intensity, from a 16S rRNA gene-based survey of farmland across North America. We then analyzed trends according to bacterial life-history frameworks and an environment-wide association survey (EWAS) to gain ecological insights. Life-history traits were assessed using genomic traits inferred from taxonomic classifications and included: genome size, *rrn* copy number, and coding density. An EWAS was conducted using 89 studies of agricultural land management. Most bacterial indicators were positively correlated with biological measures and negatively correlated with physical and chemical measures of soil health, revealing broad differences in the way management shapes bacterial associations with soil health. High soil health ratings corresponded with life-history traits associated with metabolic dependency (smaller genome and lower coding density), while lower health scores corresponded with traits selected for by environmental instability and disturbance (larger genome and multiple *rrn*). Trade-offs in community-weighted genome size explained most variation in overall health score. EWAS confirmed the importance of disturbance-adapted bacterial indicators, underscoring the impacts of tillage on soil bacterial communities. These findings provide insights into the ecological relationships between bacterial indicators and soil health and illustrate new approaches for interpreting patterns in microbiome data.

## Introduction

Managing soil health promotes the long-term fertility and ecological integrity of agricultural lands [1, 2]. Soil health encompasses a range of soil properties that contribute value to agroecosystems, including nutrient and water cycling, biodiversity, plant pathogen suppression, and pollution mitigation. Soil health is monitored using biological, physical, and chemical indicators that correspond with these functions [3–5]. Ideally, indicators should be interpretable and exhibit a dynamic response to management practices [6–9]. The soil microbiome has considerable potential to serve in this capacity. Microbial communities are highly sensitive to management practices [10–13], including those that shape soil health in agricultural systems [14–18]. The broad ecological and functional diversity of bacteria in soil provides rich information about soil conditions, which was recently used to predict soil health status [19]. However, our ability to interpret the responses of bacterial indicators is limited by our sparse understanding of the ecology and function of most bacteria in soil. Developing ways to bridge this gap will expand the utility of microbiome data in soil health monitoring.

Life-history frameworks offer a way to derive ecological information from trends in the soil bacterial communities and have provided insight into processes like carbon and nitrogen cycling [20–23]. The life-history strategies of every organism are shaped by tradeoffs between growth, survival, and reproduction, and can manifest in diverse physiological traits related to growth rate, stress tolerance, dormancy, dispersal, and resource acquisition. Life-history frameworks group organisms with similar life-history strategies into ecological categories to help interpret broad changes in microbiome structure and function. The copiotroph-oligotroph spectrum was one of the first life-history frameworks applied to the soil microbiome, grouping organisms by adaptations to high versus low resource niches with a focus on traits related to growth rate [24]. Other life-history frameworks focus on tradeoffs in the metabolic breadth between generalists and specialists [25], or a tripartite relationship between adaptations for yield, competition, and stress tolerance [23, 26, 27], or the metabolic dependency of organisms ranging from more independent (‘prototrophic’) to more dependent (‘auxotrophic’) [28]. Life-history strategies can be identified from their influences on the evolution of bacterial genomes and can be inferred from microbiome amplicon sequencing data based on phylogenetic relatedness to genomes available in public databases [29–32]. Life-history frameworks have been used to interpret changes in bacterial communities in relation to natural processes and management practices that affect soil health, such as aggregate formation [17] and cover crop and tillage practices [29, 33], respectively.

While promising, life-history frameworks remain untested in their capacity to represent the breadth of trait diversity of soil bacteria. Furthermore, many of the most active and abundant microorganisms in agricultural soils lack representative genomes, from which traits might be predicted, and are known exclusively by their phylogenetic gene markers [34–37]. Ecological information can still be derived for these organisms by profiling their phylogenetic gene markers across the growing number of publicly available sequencing projects [38, 39]. An ‘environment-wide association survey’ (EWAS) approach follows the principle of reverse ecology, where ecological attributes are inferred from the changes in abundance and distribution of a gene or genome across habitat, niche, or experiment [40]. Traditional approaches assign a trait using curated databases [41, 42], which exclude uncultured or poorly characterized taxa. This is problematic as these unclassified taxa are commonly indicative of properties relevant to soil health [19, 37, 43, 44]. In contrast, EWAS requires no prior knowledge, with information gained for any organism represented in sequencing databases [45–48]. An EWAS approach is primarily limited by the poor quality of metadata reported for most sequencing projects [49] and a historical lack of standardization in sequencing workflows. These drawbacks are partially compensated for by the sheer volume of available sequencing projects. Renewed efforts to systematize data publishing will improve the efficacy of EWAS over time [50].

This study identifies and seeks to explain the relationships between bacterial indicators of twelve conventional measures of soil health using a large survey of farmland from across North America. Our first objective was to identify the bacterial indicators of biological, physical, and chemical measures of soil health, as well as tillage practice, using 16S rRNA gene amplicon sequencing data. Our second objective was to evaluate trends in bacterial indicators using life-history frameworks and an EWAS to better understand the ecological basis for their associations with soil health. We tested whether trends in community-weighted genomic traits, corresponding to life-history frameworks, matched underlying assumptions about soil health (Table 1). Next, we obtained the environment-wide associations of bacterial indicators in a survey of 89 studies of agricultural soils and tested whether associations, grouped by study factors into broad (disturbance, management, and rhizosphere association) and specific associations (fertilization, land-use, tillage, drought etc.) were correlated with soil health. These approaches provided ecological information about the bacterial indicators of soil health, affording new perspectives on soil health management.

**Table 1.** An overview of genomic traits used to infer life-history strategies, their corresponding frameworks and the expected relationship between genomic traits and soil health. Genomic traits were derived from a collection of isolate genomes, single cell amplified genomes and metagenome-assembled genomes, assigned to bacterial taxa and used to calculate community-weighted trait averages using relative abundance information. Community-weighted averages were correlated with health ratings to test whether trends in life-history frameworks corresponded with differences in soil health. Biosynthetic gene clusters encode pathways for the production of secondary metabolites commonly involved in antibiosis and resource acquisition, which are associated with a more independent life-history strategy. CRISPR arrays corresponds to the number of arrays in a genome and not the length or number of protospacer adjacent motifs.

| Genomic trait | Life-history framework | Expected Relationship to Soil Health | Expectation | References |
| --- | --- | --- | --- | --- |
| Genome size | Generalist / competitor (large) - specialist / yield-adapted (small) | Positive | Generalists / competitors will be favored by higher quality and quantity of OM in healthier soils based on greater metabolic versatility. | Schmidt et al., 2018; Malik et al., 2020 |
| *rrn* copy number (growth rate) | Copiotroph (high) - oligotroph (low) | Negative | Copiotrophs will be favored by management practices that disturb soil and will correspond with lower health. | Roller et al., 2016; Westoby et al., 2021; Schmidt et al., 2018 |
| * Biosynthetic gene clusters | Independent (high) - dependent (low) | Negative | Independent bacteria will be prevalent in degraded soils where self-sufficiency provides greater fitness. | Wilhelm et al., 2021 |
| Coding density | - | Exploratory | Low c.d. is common in obligate mutualists, which will be more abundant in healthier soils where biomass and cell densities are highest. | McCutcheon et al., 2011; Gil and Latorre, 2012; Land et al., 2015 |
| CRISPR array | - | Exploratory | Phage abundance is positively correlated with biomass and organic matter, but the fitness tradeoffs between CRISPR and other phage defense mechanisms make this relationship difficult to predict. | Westra et al. 2019; Williamson et al., 2005; Williamson et al., 2013; Liang et al., 2019 |

## Methods

### Bacterial community and soil health data collection

Our primary dataset consisted of 778 soil samples sourced from farmland across the USA as part of an initiative by Cornell University and the USDA Natural Resources Conservation Service to characterize soil health. This dataset was used in a separate study to test the accuracy of microbiome-based machine learning for predicting soil health [19]. Our study aims to identify indicators and explore their underlying ecological basis for their association with soil health, which have yet to be examined.

Samples originated from 162 unique sites (> 1 km radius apart) termed ‘geo groups’ (n_median_ = 2 sample per group, n_mean_ = 5, n_max_ = 48). Soil health data was collected for each sample using the Comprehensive Assessment of Soil Health (CASH) framework (Table S1), which uses biological (soil organic matter, respiration, ACE protein, and active carbon, also known as ‘permanganate oxidizable organic carbon’), chemical (pH, phosphorus, potassium, and minor elements), and physical ratings of soil health (aggregate stability, available water capacity, soil texture, and surface and subsurface hardness) [7]. Data on tillage was collected for most soils (n = 599) and was coded as ‘till’ versus ‘no till.’ Higher surface and subsurface hardness ratings were inverted so that more compacted soils corresponded with higher ratings (opposite of CASH); these ratings present for a subset of samples (n = 309 and 292, respectively). Measurements for each soil property were transformed using a scoring function [7] to create a normalized rating that accounts for differences in soil texture. A total health score was then calculated from the unweighted mean of all twelve ratings.

Total DNA was extracted from soils to determine bacterial community structure and as a measure of microbial biomass [51]. DNA was extracted using the DNeasy PowerSoil Kit, as per manufacturers recommendation (QIAGEN, Germantown, MD, USA). DNA concentration was quantified using the Quant-iT™ PicoGreen™ dsDNA Assay Kit (Thermo Fisher Scientific, Inc., Waltham, MA, USA). Bacterial community composition was determined through amplicon sequencing of the V4 region of the 16S rRNA gene using Illumina MiSeq (2×250 paired-end) and dual-indexed barcoded primers (515f/806r) as previously described [19]. Operational taxonomic units (OTUs) were defined as amplicon sequence variants and assigned taxonomic classifications using *QIIME2* (v. 2020.2) [52] with dependencies on *DADA2* [38] and the *Silva* database (nr_v132) [53], respectively. Raw sequencing data was archived at the National Centre for Biotechnology Information (BioProject: PRJEB35975).

### Identifying bacterial indicators of soil health

Bacterial OTUs indicative of soil health were determined by Spearman rank correlations using the ‘rcorr’ function in the R package *Hmisc* (v. 1.34.0) [54]. Prior to correlation analyses, data was sparsity-filtered and normalized by sequencing library depth and reported as counts per thousand reads as previously described [19]. P-values were adjusted according to the Benjamini and Hochberg false discovery rate [55]. Weak correlations (r < |0.3| and *p*_adj_ > 0.05) were removed [56]. Indicator species analyses was used to identify indicators for tillage intensity using the “multipart” function in the R package *indicspecies* [57]. All analyses can be reproduced with scripts included in the Supplementary Data package.

### Community-weighted genomic traits and life-history framework analyses

OTUs were assigned trait values corresponding to genomic traits associated with life-history frameworks (Table 1). The genomic traits included: genome size and coding density (total length of coding regions / genome size) and the average abundance of *rrn* operons, CRISPR arrays and biosynthetic gene clusters per genome. Genome data was downloaded from the IMG-ER (downloaded March 15^th^, 2020) [58] and was comprised of data from isolate (n = 68,600), single-cell amplified (n = 3,400) and metagenome-assembled genomes (n = 8,800). Gene abundances were first normalized by genome size. OTUs were assigned a trait value iteratively based on taxonomic classification. Unclassified OTUs at the rank genus were progressively matched to averaged trait values at higher taxonomic ranks. Most OTUs were assigned a trait value (20,148 / 21,463; 94%) and the majority were assigned at their lowest classified taxonomic rank (58%). The community-weighted average trait values were calculated for whole bacterial communities using the weighted mean based on the relative abundance of each OTU in the community. Additionally, community-weighted *rrn* abundance was re-calculated using output from the *rrnDB* (v. 5.6) [59], yielding results consistent with calculations derived from IMG-ER data.

### Environment-wide Association Survey

We compiled 89 studies from agricultural or related terrestrial environments, totaling 14,780 individual 16S rRNA gene amplicon libraries, termed the ‘AgroEcoDB.’ Amplicon libraries associated with NCBI BioProjects with taxonomic IDs for “soil metagenome” (taxID: 410658), “compost metagenome” (702656), “decomposition metagenome” (1897463), “fertilizer metagenome” (1765030), “manure metagenome” (1792145), “rhizosphere metagenome” (939928), and “wood decay metagenome” (1593443) were downloaded on May 15^th^, 2020. The final database was filtered from an initial 729 BioProjects to 89 based on the following criteria: (i) common overlap of the V4 region of the 16S rRNA gene, (ii) experimental manipulations, when used, were typical of agricultural management, and (iii) contained at least 15 samples with well-curated metadata (details in Supplementary Methods). Study factors were categorized by management categories (e.g., inorganic versus organic fertilizer and other broad strategies, like crop rotation), disturbance (tillage, drought etc.), plant association (bulk vs. rhizosphere soil), biome (grassland vs. cropland) and other minor categories (decomposition, soil depth etc.) (Table S2). Indicator species analyses was used to attribute an indicator value to OTUs in the AgroEcoDB based on each individual study factor. Study factors were coded to reflect expectations about their relationship with soil health and indicator values were scored as positive or negative based on whether the relationship was positively (reduced tillage, OM management, etc.) or negatively associated with soil health (Table S2). Our subsequent analyses were a test of whether these assumptions were supported by trends in soil health data.

### Bioinformatic Analyses

Statistics were performed using R (v. 4.0.3) [60] with dependency on the following packages: *reshape2* (v. 1.4.4), *ggplot2* (v. 3.3.2), *plyr* (v. 1.8.6) [61–63], and *phyloseq* (v. 1.34.0) [64]. Observed species richness and Shannon diversity estimates were averaged from the ‘plot_richness’ function based on 200 permutations of rarefied counts (n = 1,000 reads). Permutational multivariate analysis of variance (PERMANOVA) was performed on Bray-Curtis dissimilarity using the R package *vegan* (v. 2.5.7) [73] with 999 permutations. PERMANOVA was repeated with 50 permutations of factor order to obtain average R^2^ values. The relative importance of community-weighted traits and environment-wide associations for explaining variation in community composition was compared with *relaimpo* [65]. Co-occurrence networks were constructed for bacterial taxa (aggregated by genus) based on whether two genera shared a common indicator status for each of the twelve soil health ratings. Edges were weighted by the number of OTUs co-occurring between nodes (i.e., each genus). Indicators with negative and positive correlations with ratings were visualized in separate networks using Gephi (v.0.9.2) [66] with network topography determined by the Yifan Hu ‘proportional’ force-directed graphing algorithm (relative strength = 2) [67].

## Results

### Relationships among measures of soil health

Biological ratings were highly interrelated and positively correlated with total health score and with aggregate stability, a physical property influenced by biological activity (Figure 1; full matrix in Figure S1). Total health score was negatively correlated with surface and subsurface hardness ratings (where a higher rating indicates greater compaction), tillage intensity, and sand content. DNA yield was significantly positively correlated with soil health (r = 0.51; *p* < 0.001), but was influenced by clay content, likely as a result of higher sorption reducing DNA extraction efficiency (Figure S2). Bacterial indicators of soil health ratings were apparent despite broad variation in community structure due to geography (PERMANOVA; R^2^ = 0.58), followed by tillage intensity (0.02), soil texture class (0.009), pH (0.008) and total health score (0.006; Table S3).

**Figure 1.**
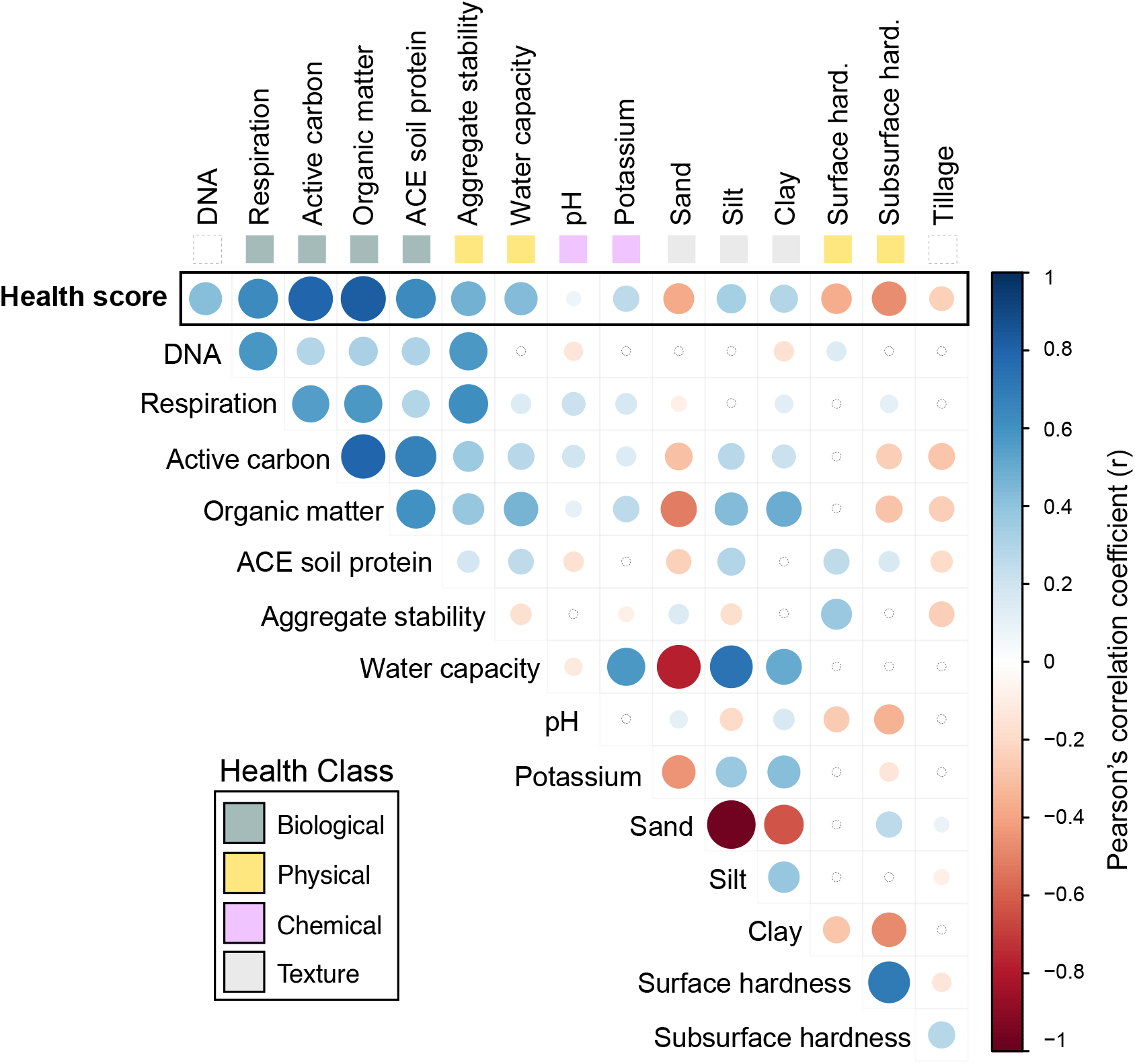
A correlation matrix showing the relationships among soil health ratings, DNA yield and soil texture. Health ratings are grouped by biological (green), physical (yellow), texture (gray), and chemical (pink) classes. The strength of each Pearson’s correlation corresponds with the intensity of color of blue (r > 0) and red (r < 0). The larger the circles the lower the *p*-value with non-significant correlations indicated by a small, colorless circle. Higher hardness ratings correspond with more compaction which explains the negative correlation with total health score. Minor elements and phosphorus ratings had few significant correlations and are not shown here – a complete matrix is provided as Figure S1.

### Bacterial indicators of soil health

OTUs were correlated with each of the twelve health ratings and total health score to identify bacterial indicators (r > | 0.3 | and *p*_adj_ < 0.05). A subset of OTUs (1,874 / 21,463) were identified as indicators of one or more health ratings (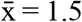 ratings per OTU; max = 5). Indicators were taxonomically diverse (348 different classifications at rank genus) with most belonging to either a candidate group or were unclassified below the rank of family (62%). Approximately twice as many unclassified or candidate genera (1.9-fold) were present in the indicator set (215 / 348) compared to the overall dataset (430 / 943). Positively correlated OTUs were primarily indicative of biological ratings, while negatively correlated OTUs were indicative of physical or chemical ratings (Figure 2). Many genera (46%) contained a mix of indicator OTUs responding either positively or negatively to different health ratings. However, the majority of indicator OTUs were correlated in a consistent direction with one or more health ratings (96%; 1,798 / 1,874).

**Figure 2.**
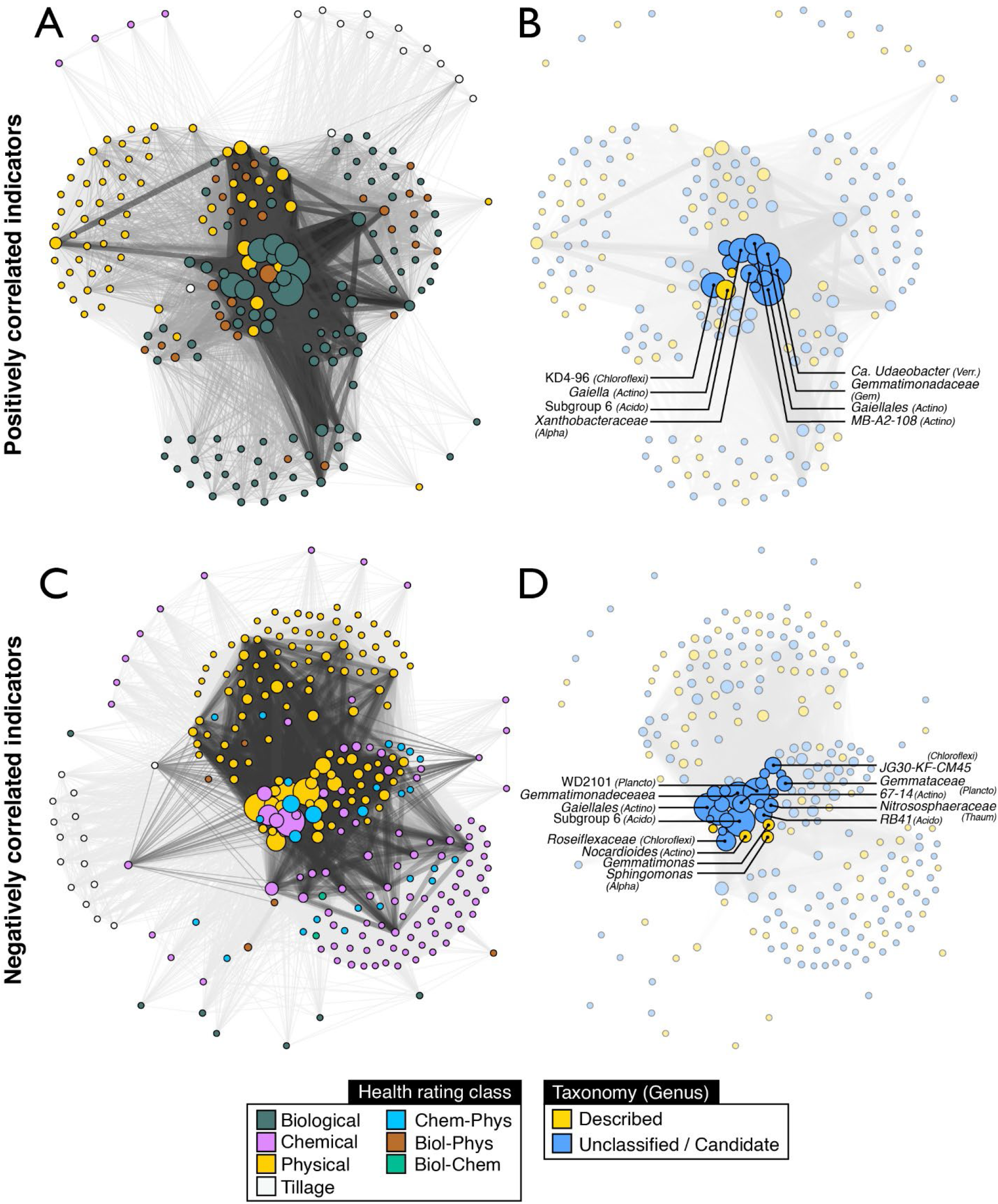
Network diagrams displaying the co-occurrence of indicators among the three classes of health ratings. Networks were divided based on whether indicators exhibited positive (a and b) or negative correlations (c and d) with health ratings to illustrate the broad differences between indicators of biological versus chemical or physical ratings. Nodes represent indicator OTUs aggregated according to their lowest resolved taxonomic rank (scaled by the total number of OTUs), while edges represent co-occurrence of any OTU as an indicator for the same health rating. Edge weights are scaled by the number of co-occurring OTU common between nodes. In (a) and (c), nodes are colored according to health rating class and, in (b) and (d), according to whether a taxon is represented by a described species. In (a) and (c), nodes were colored based on majority rules according to the number of OTUs representing a given health class. Classes were hyphenated when no majority was achieved.

The main indicators of high biological health status were members of unclassified KD4-96 (*Chloroflexi*), Candidatus *Udaeobacter* (*Verrucomicrobia*), unclassified *Xanthobacteraceae* (*Alphaproteobacteria*) and unclassified MB-A2-108 and *Illumatobacteraceae* (*Actinobacteria*; full list in Table S4). The most consistent indicators of poorer physical, chemical and total health score were classified as JG30-KF-CM45 (*Chloroflexia*), Ca. *Nitrososphaeraceae* (*Archaea*; *Thaumarchaeota*), *Sphingomonas* (*Alphaproteobacteria*), and RB41 (*Acidobacteria*; *Pyrinomonadaceae*). Many taxa occurring at high relative abundances (1-5% of total read counts) were differentially abundant in tilled soils, with most occurring at increased relative abundances in tilled fields (n_OTU_ = 292) in contrast to no till systems (n_OTU_ = 18). The predominant indicators of tilled soils were members of *Alphaproteobacteria* (*Sphingomonadaceae*, *Rhizobiaceae* and *Caulobacteraceae*) *Pyrinomonadaceae*, *Chthoniobacter* (*Verrucomicrobia*) and *Terrabacter* (*Actinobacteria*). The main indicators of untilled fields were the aforementioned indicators of high biological health ratings, as well as *Gaiella* (*Actino*.) and unclassified *Solirubrobacterales* (*Actino*.).

### Life-history frameworks

We evaluated trends in the community-weighted averages of genomic traits represented in several life-history frameworks (Table 1). Communities with a higher prevalence of bacteria with larger genomes and those which encoded more biosynthetic gene clusters (BGCs) were associated with lower total health score (Figure 3A), traits which correspond with more competitive, independent taxa. Communities with a higher prevalence of bacteria with high coding density were associated with higher health score, a trait associated with more dependent taxa. The relationships among traits in community-weighted data partially reflected the existing relationships observed in genomic data (Mantel statistic r = 0.64; *p* = 0.01). However, the relationships among genome size, BGCs, and *rrn* copies differed from these existing relationship in community-weighted data (Figure 3B).

**Figure 3.**
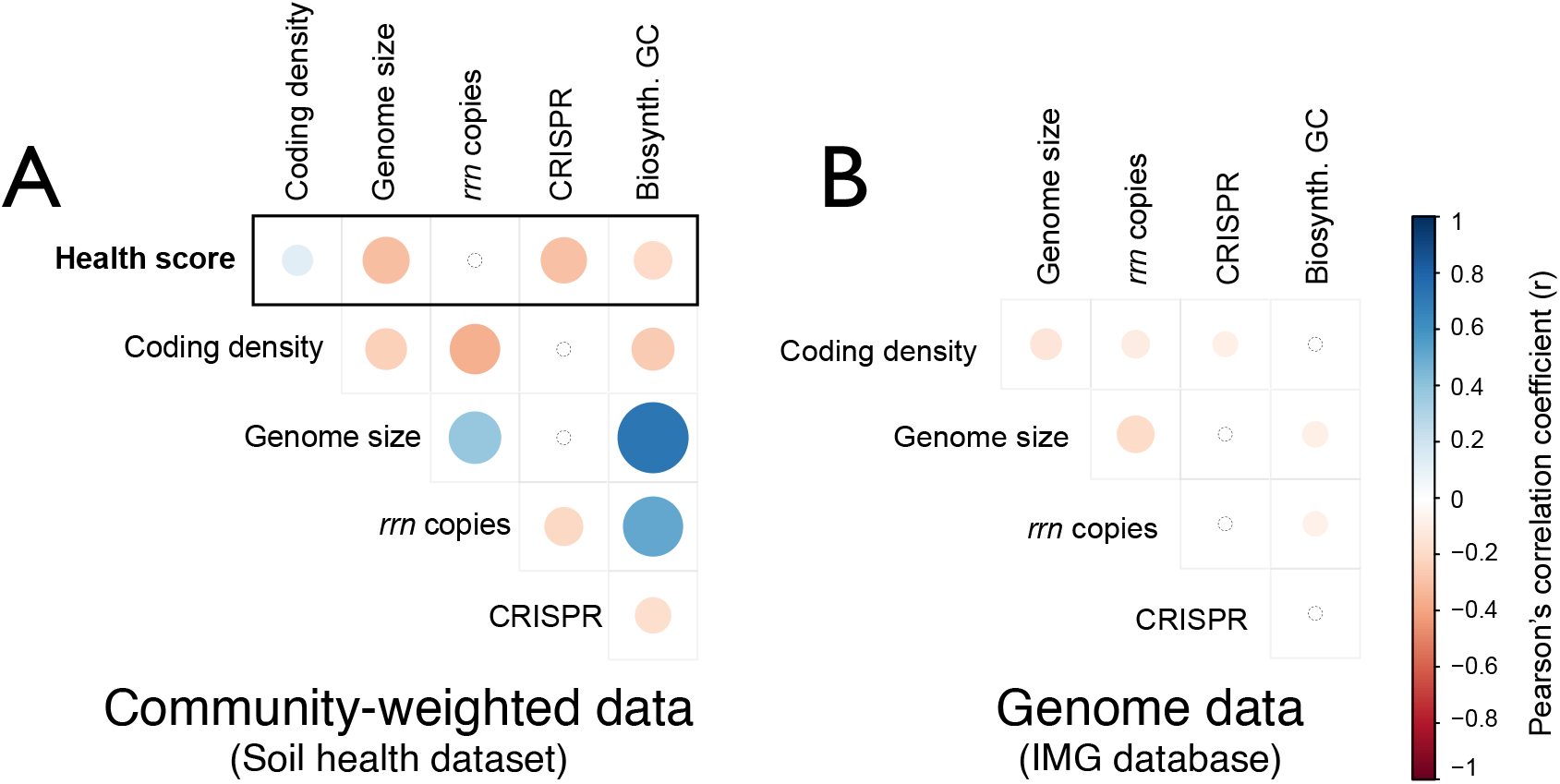
Correlation matrices illustrating the relationships among genomic traits and total health score based on (a) average trait scores weighted by the relative abundance of taxa-specific traits values (i.e., community-weighted data) in the soil health data or (b) the genomic database used to assign trait values to taxa. This side-by-side comparison illustrates that the relationships among traits in community-weighted data partially reflected the existing relationships observed in the genomic data (Mantel statistic r = 0.64; *p* = 0.01). The strength of each Pearson’s correlation corresponds with the intensity of color of blue (r > 0) and red (r < 0), and only significant correlations are colored.

Community-weighted *rrn* copy number was not correlated with total health score (r = 0.003). However, community-weighted *rrn* copy number was significantly higher in tilled soils (Wilcoxon, *p* < 0.001; Figure 4) and exhibited significant correlations with soil surface and sub-surface hardness rating (Figure 5A). The trends in community-weighted *rrn* copy number were driven by the increased relative abundances of *Georgenia* (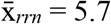; *Actino*.), *Bacillaceae* (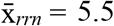; *Firmicutes*), and *Planococcaceae* (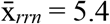; *Firmicutes*). Community-weighted genome size and BGCs abundance were negatively correlated with total health score, and like *rrn* copy number, showed increased prevalence in tilled soils (Figure 4). Genome size and BGCs were primarily correlated with biological ratings, unlike *rrn* copy number which was exclusively correlated with physical or chemical ratings (Figure 5A).

**Figure 4.**
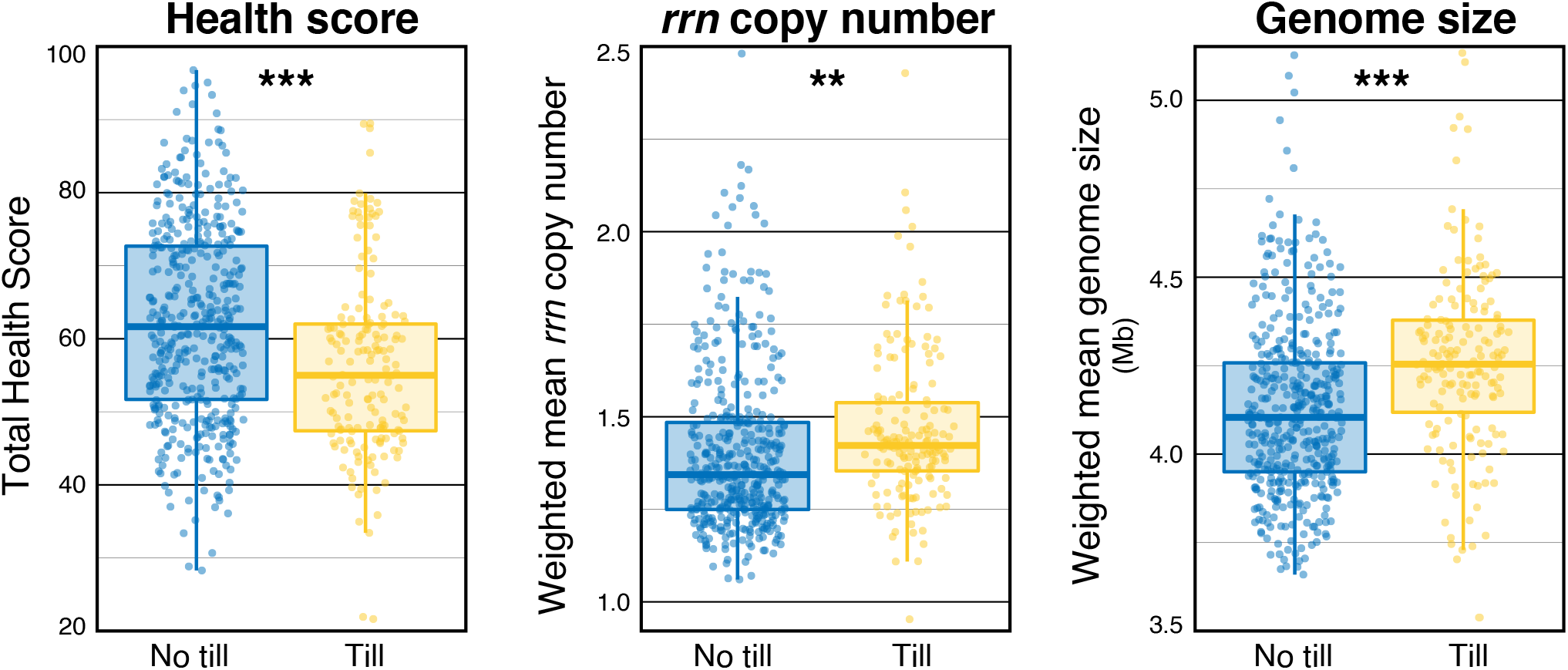
A comparison of total health score and community-weighted traits in tilled and untilled soils visualized as box and whisker plots. Significant differences are denoted with asterisk based on t-tests (** *p* < 0.01; *** *p* < 0.001).

**Figure 5.**
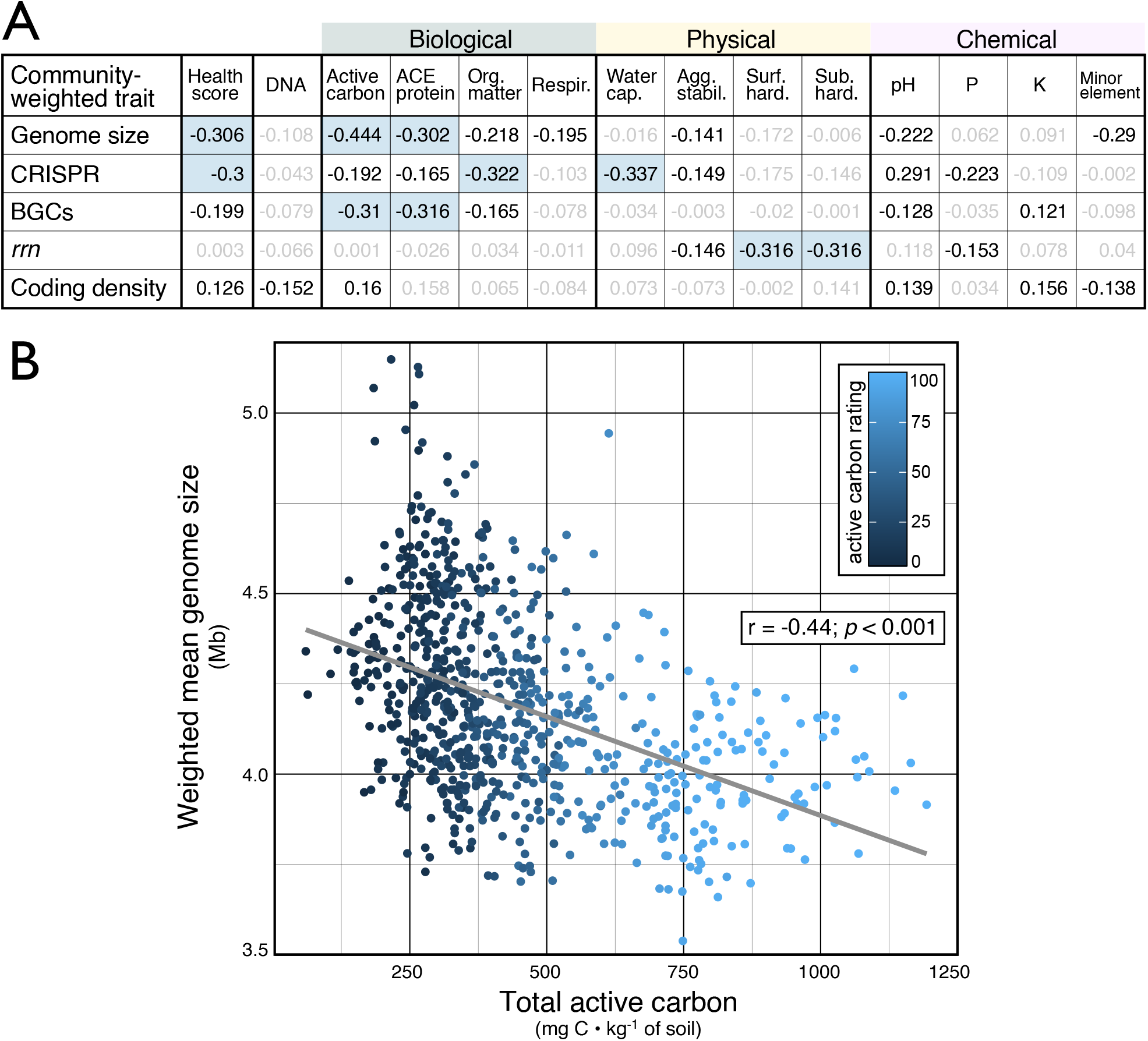
A summary of correlations between (a) soil health ratings and community-weighted traits and (b) a plot of community-weighted genome size versus active carbon content (colored by rating). In (a), all Pearson’s r > | 0.3 | are shaded blue and all significant correlations are shown in bold. In (b), the relationship was not due to biases arising from differences in the proportion of unclassified taxa assigned traits which was not correlated with active carbon rating (r = 0.06, *p* = 0.1; see Figure S7).

Community-weighted coding density was positively correlated with total health score and with ratings of OM quality (active carbon and ACE protein; Figure 5A), but negatively correlated with DNA yield, a proxy for microbial biomass. The bacterial indicators of DNA yield with the lowest coding density belonged to candidate groups of *Chloroflexi* (*Ktedonobacterales*: HSB_OF53-F07 and JG30a-KF-32; 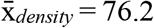) and a family of *Planctomycetes* (*Gemmataceae*; 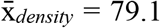). Community-weighted CRISPR array abundance exhibited some of the strongest correlations with health ratings and was negatively correlated to water capacity, measures of OM quality, and total health score (Figure 5A), and positively correlated with sand content (r = 0.44; *p* < 0.001).

### Environment-wide associations

The majority of OTUs in the soil health dataset were present in the AgroEcoDB (n_OUT_ = 17,818 / 21,573), representing a total of 96.9% of sequences. A total of 8,760 overlapping OTUs were identified as significant indicators (*p* < 0.01) of one or more study factors in the AgroEcoDB. Indicator values for these OTUs were used to calculate community-weighted averages of broad categories and sub-categories to assess general correlations between the environment-wide associations of bacteria and soil health. Community-weighted environment-wide associations explained more variation in bacterial community composition than community-weighted genomic traits (Table 2A), primarily based on variation captured by associations with disturbance and management strategy. However, community-weighted genome size explained the most variation in total health score (Table 2B). Community-weighted environment-wide associations with disturbance (r = 0.12; *p* = 0.001) and management (r = 0.11; *p* = 0.002) were positively correlated with total health score, demonstrating that our coding of study factors (i.e., indicators of health-promoting factors were assigned a positive indicator value) generally conformed with expectations about soil health.

**Table 2.** Comparisons of the variation in (a) community composition and (b) total health score explained by community-weighted traits and environment-wide associations (shaded) according to R^2^ values from PERMANOVA, in (a), and the relative importance of each variable in linear regression (RELAIMPO) in (b). Community composition was compared based on Bray-Curtis dissimilarity. Each panel contains two separate analyses – one based on the broad categorization of environment-wide associations (upper) and another based on sub-categorization of study factors (see Table S2 for details).

| A | Variation in Community Composition |  |
| --- | --- | --- |
|  | Community-weighted trait / environment-wide association | R<sup>2</sup> |
| Category | Disturbance | 0.045 |
|  | Management | 0.039 |
|  | Genome size | 0.035 |
|  | Plant association | 0.03 |
|  | Coding density | 0.019 |
|  | *rrn* copy number | 0.017 |
| Sub-category | Organic matter quality | 0.031 |
|  | Tillage | 0.03 |
|  | Organic vs. conventional | 0.03 |
|  | Genome size | 0.03 |
|  | Drought | 0.022 |
|  | Land use | 0.022 |
|  | Host association | 0.021 |
|  | Coding density | 0.017 |
|  | *rrn* copy number | 0.016 |
|  | NPK fertilization | 0.013 |
| Community-weighted trait |  |  |
| Environment-wide association |  |  |

| B | Variation in Soil Health Score |  |
| --- | --- | --- |
|  | Community-weighted trait / environment-wide association | Rel. Imp. |
|  | Genome size | 71 |
|  | Management | 9.7 |
|  | Coding density | 7.2 |
|  | *rrn* copy number | 5.9 |
|  | Disturbance | 5 |
|  | Plant association | 1.1 |
|  | Genome size | 36 |
|  | Tillage | 20 |
|  | Organic matter quality | 12 |
|  | Drought | 9.4 |
|  | NPK fertilization | 5.9 |
|  | Coding density | 4.7 |
|  | *rrn* copy number | 3.6 |
|  | Host association | 3.1 |
|  | Land use | 2.9 |
|  | Organic vs. conventional | 2.3 |

The environment-wide associations of the most abundant bacterial indicators driving the strong correlation between community-weighted genome size and active carbon were investigated (Figure 5B). The bacterial indicators of active carbon with the largest estimated genomes were classified as *Chthoniobacter* (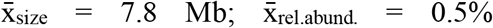 of total counts), *Geodermatophilaceae* (4.8 Mb; 1.0%), and *Sphingomonas* (4.2 Mb; 1.7%), and those with the smallest were: *Gaiella* (1.5 Mb; 2.1%), Ca. *KD4-96* (2.3 Mb; 4.9%), and Ca. *Udaeobacter* (2.7 Mb; 3.5%). Of these representative taxa, those with larger genomes had consistently higher relative abundances in tilled soils and those with low active carbon ratings, while the reverse was true for those with smaller genomes (Figure S3). According to environment-wide associations, all representative taxa were primarily associated with bulk soil rather than rhizosphere soil (Figure 6A). Their associations with soil disturbance were consistent, where taxa with larger genomes were generally positively associated with disturbance (Figure 6B). These trends were driven by disturbances related to tillage (Figure S4) and long-term irrigation regime (Figure S5). The environment-wide associations with broader management strategies (crop rotation, land-use, and fertilization) were variable (Figure 6C).

**Figure 6.**
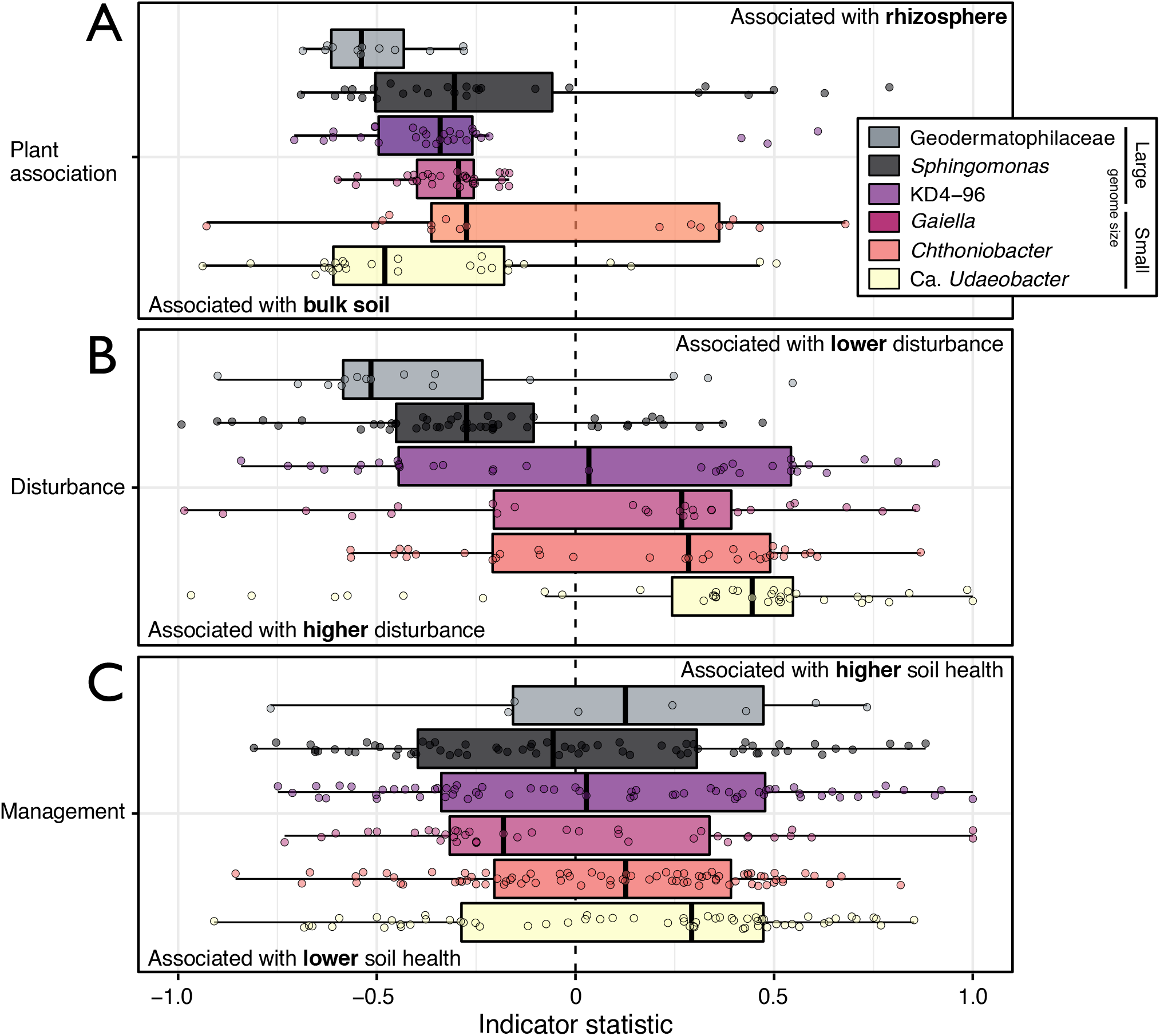
Environment-wide associations of the most abundant bacterial indicators representing three of the largest or smallest average genome sizes. Each plot shows every OTU identified as an indicator for study factors grouped by (a) plant association (i.e., bulk versus rhizosphere soil), (b) soil disturbance and (c) broad management strategy. In (b) and (c), indicator values were assigned as positive or negative based on whether a factor was a reference (i.e., no till) or treatment (till) with details of designations provided in Table S2. Only OTUs shared among the AgroEcoDB and soil health data were included. Collectively, these six taxa account for 14% of reads in amplicon libraries.

Active carbon is positively correlated with particulate forms of soil OM (POM), such as cellulose and lignin [68, 69]. Therefore, we investigated whether the representative taxa had preferences for soluble (DOM) versus particulate forms of POM in studies utilizing soil organic matter amendments in the AgroEcoDB. *Sphingomonas* were indicators of POM amendment (Indicvalue = 0.99, *p* = 0.03) in a study comparing xylose versus cellulose soil amendment [70], while no other representatives were indicators of either DOM or POM in studies comparing amendments of cellobiose versus straw (PRJNA397131), nine substrates ranging in solubility (PRJNA668741) or a wide variety of soluble and insoluble substrates (PRJNA594403). Representative taxa were more impacted by tillage than by the retention of plant residues in a study on soil management practices (Figure S4) [70], indicating responses to disturbance may play a greater role than functions in OM decomposition.

## Discussion

We sought ecological insights into the bacterial indicators of soil health by using life-history frameworks and an EWAS. Our findings lend further evidence that the abundance patterns of soil bacteria reflect changes in soil conditions relevant to soil health [18, 29] and, in our case, at a spatial scale and with an operational dataset that suggest a broad utility in monitoring. A diverse set of 348 unique bacterial genera were identified as indicators of one or more soil health rating, yet the majority (62%) were unclassified beyond the family level. This finding underscored the need for alternative strategies, like life-history frameworks and EWAS, to extend beyond our knowledge of cultivated bacteria to gain insights into bacterial indicators of soil health.

### The effects of soil disturbance on soil health according to life-history traits

The relationship between community-weighted genome size and total health score was among the strongest correlations observed for any genomic proxy for life-history traits (Figure 5) and was the most important predictor of variation in total health score (Table 2B). On average, communities with a greater proportion of bacteria with larger genomes occurred in soils with lower health score, and lower biological ratings. This was contrary to our expectation that a large genome, and the accompanied functional versatility, would provide competitors / generalists with an advantage in soils characterized by a greater abundance and diversity of organic matter (i.e., healthier soils). The strong negative correlation between community-weighted genome size and active carbon (Figure 5B), suggested the trend may be driven by the relationship of indicators with OM quality. However, the environment-wide associations of bacterial indicators of OM quality explained less variation in total health score than genome size or tillage (Table 2B). Furthermore, the main taxa driving trends in genome size did not exhibit associations with experimental amendments of DOM or POM but, instead, were affected by disturbances like tillage (Figure S4) and irrigation regimes (Figure S5).

Bacteria with larger genomes are more abundant in habitats characterized by high environmental variability, where their expanded regulatory and metabolic capabilities provide enhanced fitness [71, 72]. Thus, the larger community-weighted genome size associated with lower soil health, and tillage, indicate a higher proportion of taxa adapted to environmental instability, consistent with observations in tidal systems [73]. Conversely, bacteria with smaller genomes, associated here with higher health, tend to be marked by dependencies on services provided by microbial community members and may, therefore, be more sensitive to disruptions. For example, Ca. *Udeaobacter* possess a remarkably reduced genome [74] and were among the strongest indicators of high health rating and were most impacted by disturbances (Figure 6; Figure S3; Figure S4). Along with a reduced genome size, Ca. *Udeaobacter* exhibit high levels of auxotrophy and antibiotic tolerance [74, 75], characteristic of the life-history strategy of dependent organisms, exemplified by members of *Planctomyces* [28, 76]. Trade-offs in the putative capacity to produce antibiotics were apparent in the positive correlation between community-weighted BGC abundances and genome size. These observations suggest that life-history strategies marked by independence (large genomes and higher BGC capacity) and dependence (small genome and antibiotic tolerance) correspond with adaptations to environmental / community stability which correlate with soil health.

Our evidence suggests that community-weighted genome size may serve as an indication of the effects of management on environmental complexity and stability, integrating the myriad effect of disturbances from fertilization, tillage, watering, crop rotations, and harvesting. However, we could not determine whether community trends were driven by the growth dynamics of populations with larger or smaller genomes, or both, since sequencing data is a measure of relative abundance [77]. Disentangling this dynamic is important for understanding the exact nature of the relationship with genome size and, in future, may be addressed using internal standards [78–81].

While our analyses revealed patterns in the genomic traits and environment-wide associations of the soil microbiome, they also illustrated the challenges of categorizing the life-history strategies of bacteria into broad frameworks. For example, *Chthoniobacter* had a larger genome (ranging from 3.6 − 7.8 Mb) and, like other taxa with larger genomes, were more prevalent in tilled soils (Figure S3). However, unlike taxa with larger genomes, *Chthoniobacter* were indicative of higher health ratings, similar to their close relative Ca. *Udeaobacter* (family *Chthoniobacteraceae*) which have substantially smaller genomes (2.7 –3.2 Mb) and were less prevalent in tilled soils. Investigating the ecology of these abundant, closely related soil taxa may reveal differences in life-history strategies that illuminate more specific effects of soil management.

### Indicators of physical and chemical ratings

Bacterial indicators were primarily negatively correlated with physical and chemical ratings in contrast to largely positive correlations with biological measures (Figure 2). These overarching trends may correspond with differences in the nature of the management practices driving the associations between bacterial populations and soil health. The primarily negative correlations with physical and chemical properties may reflect the increased relative abundance of stress tolerant taxa or taxa with metabolic functions associated with nutrient regimes experienced at lower ends of the chemical and physical health spectrum (i.e., lithoautotrophic organisms or stress-tolerators). For example, the increased relative abundance of bacterial populations classified to the family *Nitrososphaeraceae* were strongly indicative of soils with poorer soil health. *Nitrososphaeraceae* are ammonia-oxidizing archaea which are commonly enriched in conventionally-managed agricultural soils fertilized with ammonia [18, 82] and exhibited environment-wide associations with fertilizer use in the AgroEcoDB (Figure S6). Conversely, the positive correlations with biological ratings may correspond with an increased relative abundance of organoheterotrophs in soils with higher OM quantity and quality, consistent with the positive correlations with respiration rating (activity) and DNA yield (biomass).

Community-weighted *rrn* copy number, a correlate of bacterial growth rate [83], was the only trait that had a significant relationship with hardness ratings and the only trait not correlated with total health score or biological ratings (Figure 4). Community-weighted *rrn* copy number was significantly higher in tilled fields, matching our expectation that disturbance would favor fast-growing copiotrophs, consistent with previous findings [84]. Notably, hardness ratings were negatively correlated with *rrn* copy number, indicating communities in more compacted soils tended to encode a higher number of *rrn* operon. This matched our expectation that higher *rrn* copy number would be associated with more degraded soils (Table 1), though the nature of the relationship between fast-growing taxa and soil compaction is unclear. These results confirm the influence of physical disturbance on bacterial life-history strategies related to growth rate and their potential to serve as indicators of soil health.

### Exploring relationships between coding density, CRISPR array abundance and soil health

As part of an exploratory analysis, we calculated community-weighted coding density as a proxy for the prevalence of metabolic dependency. Low coding density is a genomic signature of obligate forms of dependency apparent in epibionts, endobionts and parasites, resulting from the relaxation of selection pressures and an accumulation of pseudogenes [85, 86]. We expected lower community-weighted coding density may indicate higher trophic dependency, which we expected in more developed, healthier soils. The inverse relationship between community-weighted coding density and DNA yield supported this expectation, showing that soils with more biomass tended to have higher representation of putatively dependent taxa with lower coding density. OTUs belonging to the family *Gemmataceae* 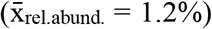 were indicators of high DNA yield and exhibited one of the lowest predicted coding density 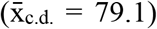. Prior evidence shows that *Gemmataceae* are dependent on nutrient supplementation from hosts or community members [76]. However, contrary to expectations, overall community-weighted coding density was positively correlated with total health score. The interpretation of this trend is complicated by our limited understanding of the full scope of evolutionary forces shaping coding density.

We explored the relationship between community-weighted CRISPR array abundance and soil health as a proxy for bacteriophage pressure. CRISPR array abundance was the only genomic trait to exhibit strong inverse correlations with available water capacity and OM ratings, and to have a positive correlation with sand content. These observations are consistent with the observation that CRISPR provides greater fitness in low resource conditions relative to other phage defense mechanisms [87]. However, our observations run counter to expectations that community-weighted CRISPR array abundance would be greatest with wet, OM-rich soils, where phage abundances are typically highest [88, 89]. This relationship requires further study, especially since CRISPR array abundance is a measure of the number of arrays, not the total length or number of protospacers, which may better correlate with phage exposure [90].

## Conclusions

Life-history frameworks and environment-wide association surveys provide new ways to interpret and explore trends in microbiome data. Both approaches proved useful for studying relationships between the soil microbiome and soil health, indicating their general utility in the study of other diverse and poorly characterized environmental microbiomes. Several community-weighted genomic traits differed across the spectrum of soil health, and our EWAS provided a means to test hypotheses about poorly described, but important, indicators of soil health. With these methods, we found evidence that bacterial indicators of soil health largely corresponded to disturbance-sensitive and stress-tolerant populations. Future research is needed to determine whether disturbance-adapted bacteria not only reflect soil health but affect processes related to healthier soils. Particular attention should be paid to their relationship with OM quantity and quality, since a larger genome has been linked to lower carbon use efficiency [91]. As such, our data suggests soil with a lower health status may be prone to carbon loss through respiration. Overall, our study illustrates the value in considering the ecological attributes of bacterial indicators of soil health to better understand and evaluate the effects of management practices.

## Acknowledgements

We thank Christopher DeRito at Cornell University for his assistance in preparing amplicon sequencing libraries. This work was supported by the USDA National Institute of Food and Agriculture, Hatch project 1010520; by the U.S. Department of Energy, Office of Biological & Environmental Research Genomic Science Program under award number DE-SC0016364; and by the USDA Natural Resource Conservation Service Conservation Innovation Grant, agreement number: 69-3A75-16-039. Any opinions, findings, conclusions, or recommendations expressed in this publication are those of the author(s) and do not necessarily reflect the view of the National Institute of Food and Agriculture (NIFA), the United States Department of Agriculture (USDA), or the United States Department of Energy.

## Author Contributions

RCW performed all data analysis, research, and writing. JPA, KSMK and HMV managed sample collection and soil health testing. DHB guided all research efforts, including analyses and writing.

---

**Figure S1.**
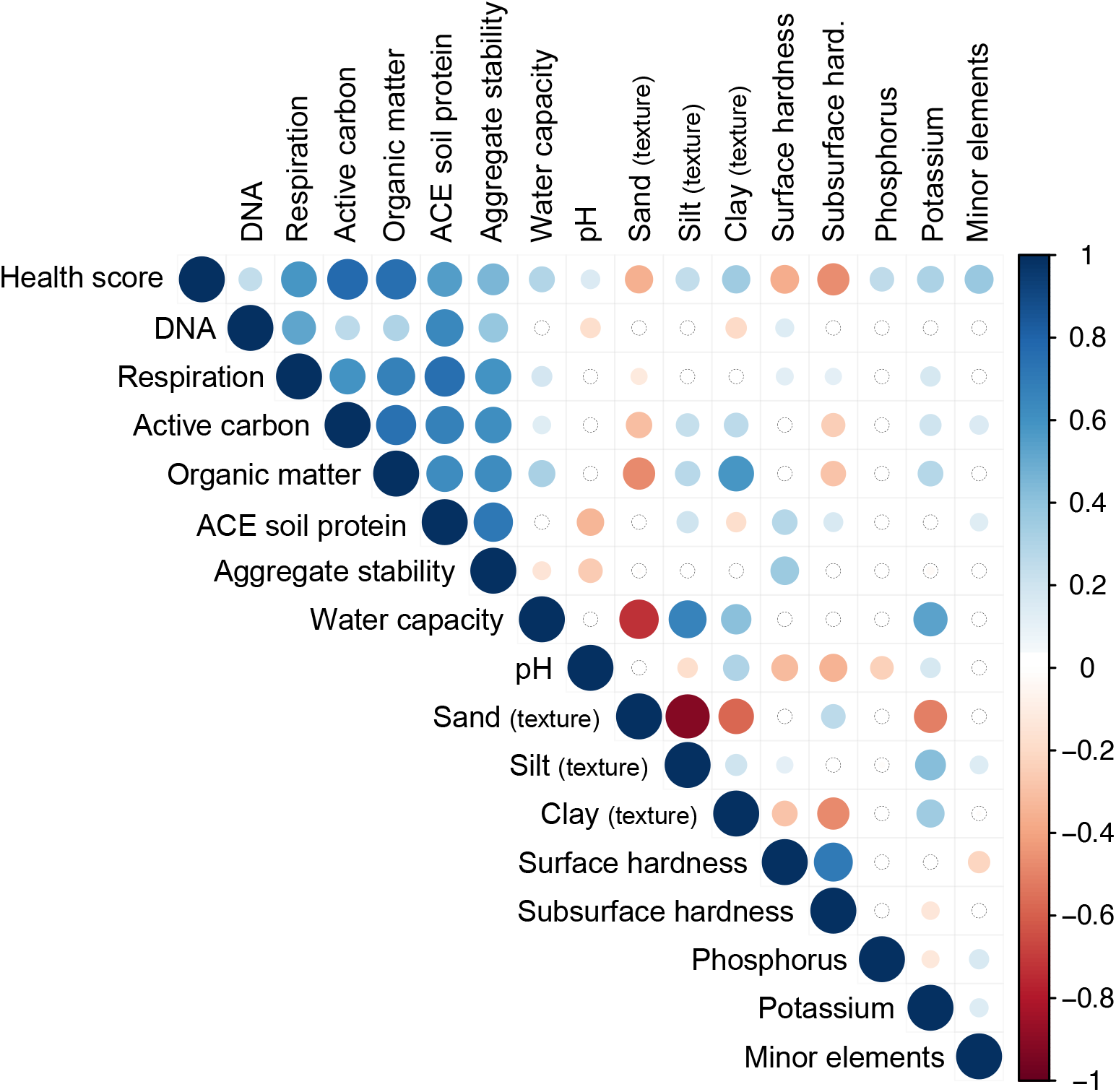
Correlation plot showing pairwise Pearson’s r values for all soil health metrics. Non-signifcant correlations are depicted by a hollow circle.

**Figure S2.**
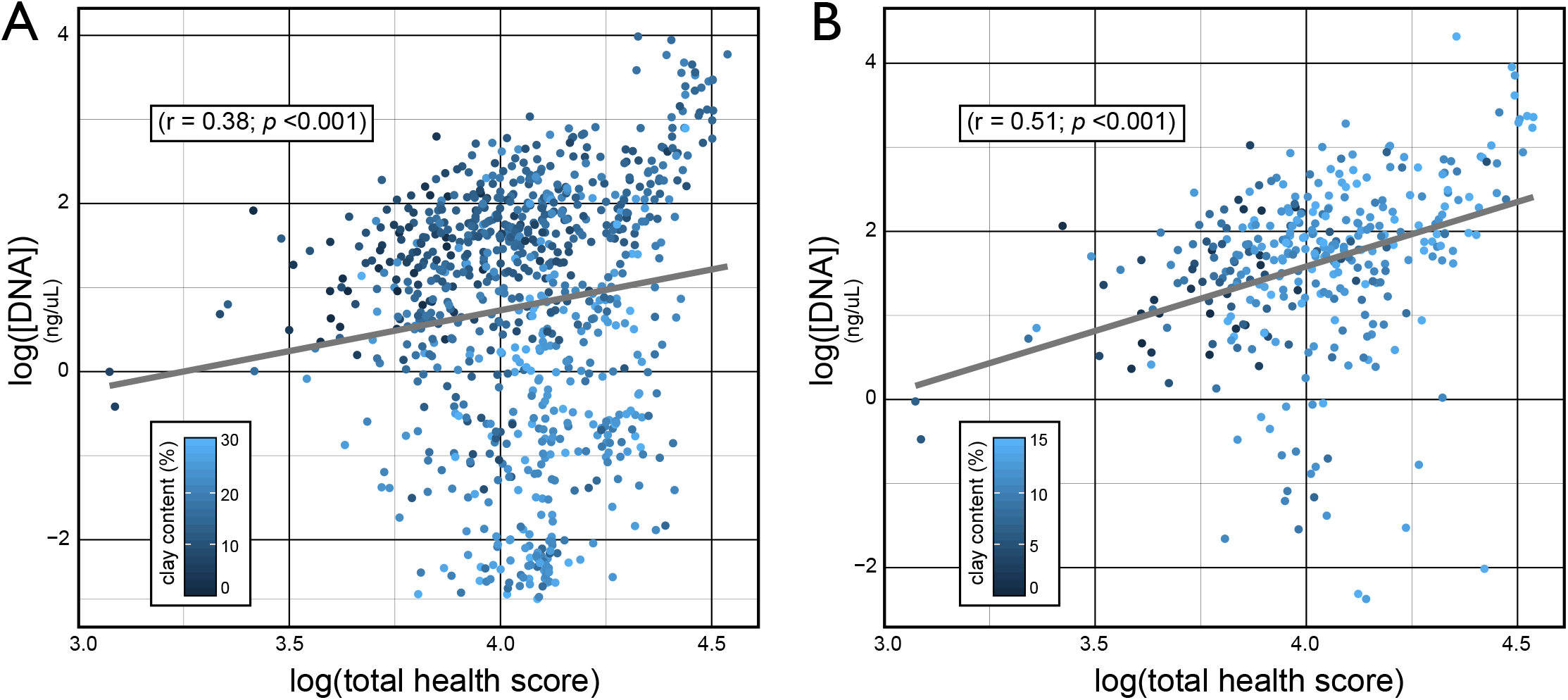
The correlation between soil DNA yield and total soil health rating was confounded by high clay content (A) and was improved following the exclusion of soils with > 15% clay content (B).

**Figure S3.**
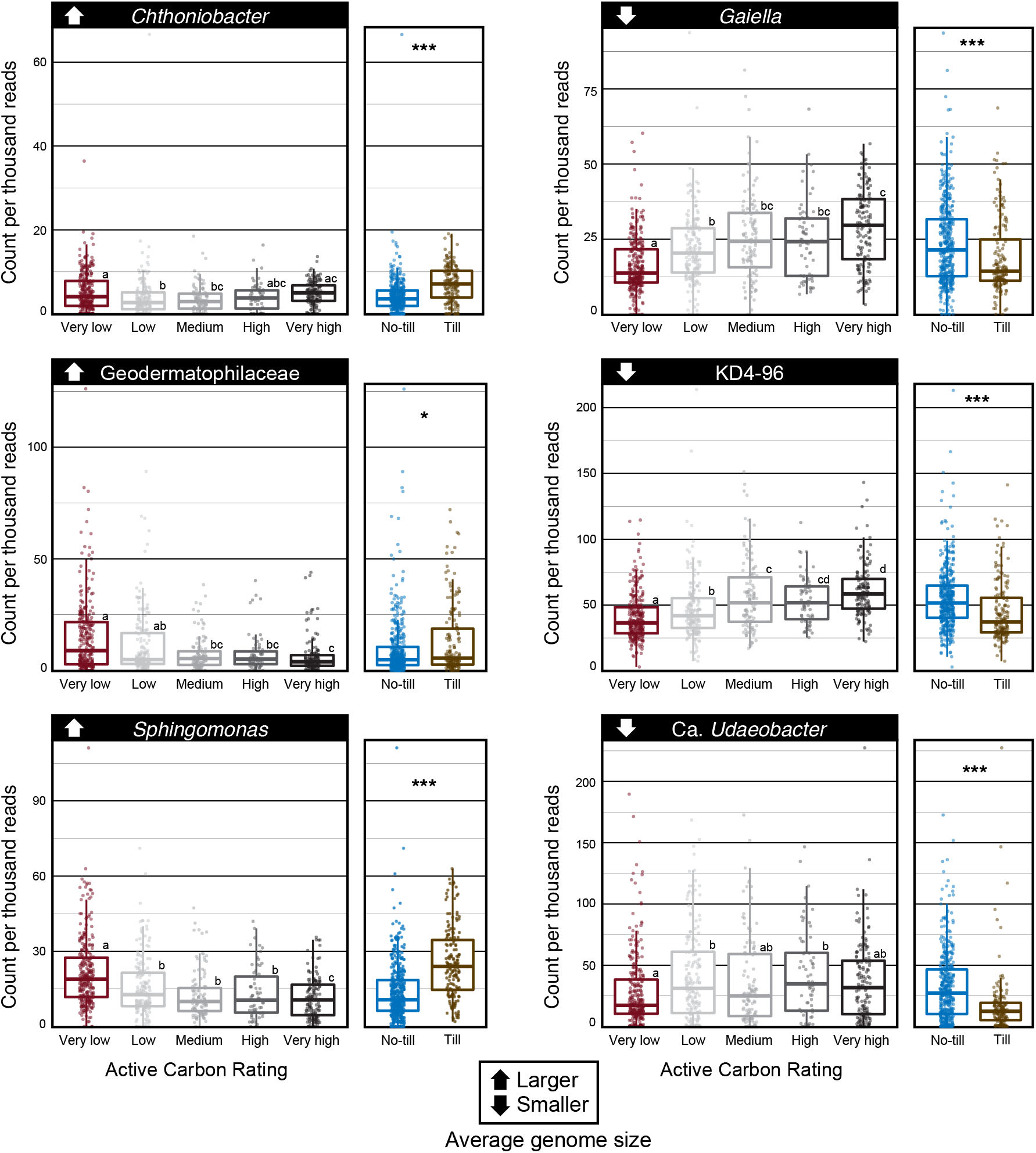
The relative abundance of six bacterial indicator taxa according to active carbon rating and tillage status. On the left, the three indicators with the largest genomes, exhibiting higher relative abundances at lower active carbon ratings and in tilled soils. On the right, the three indicators with the smallest genomes, which exhibited the opposite trends. Collectively, these taxa comprised 14% of all reads and contributed to trends shown in Figure 4 and Figure 5B. Active carbon ratings were divided into categories based on ranges of 20, from very low [0 − 20) to very high (80-100]. Pairwise statistically significant differences (p < 0.05), according to Tukey HSD, are denoted by lettering.

**Figure S4.**
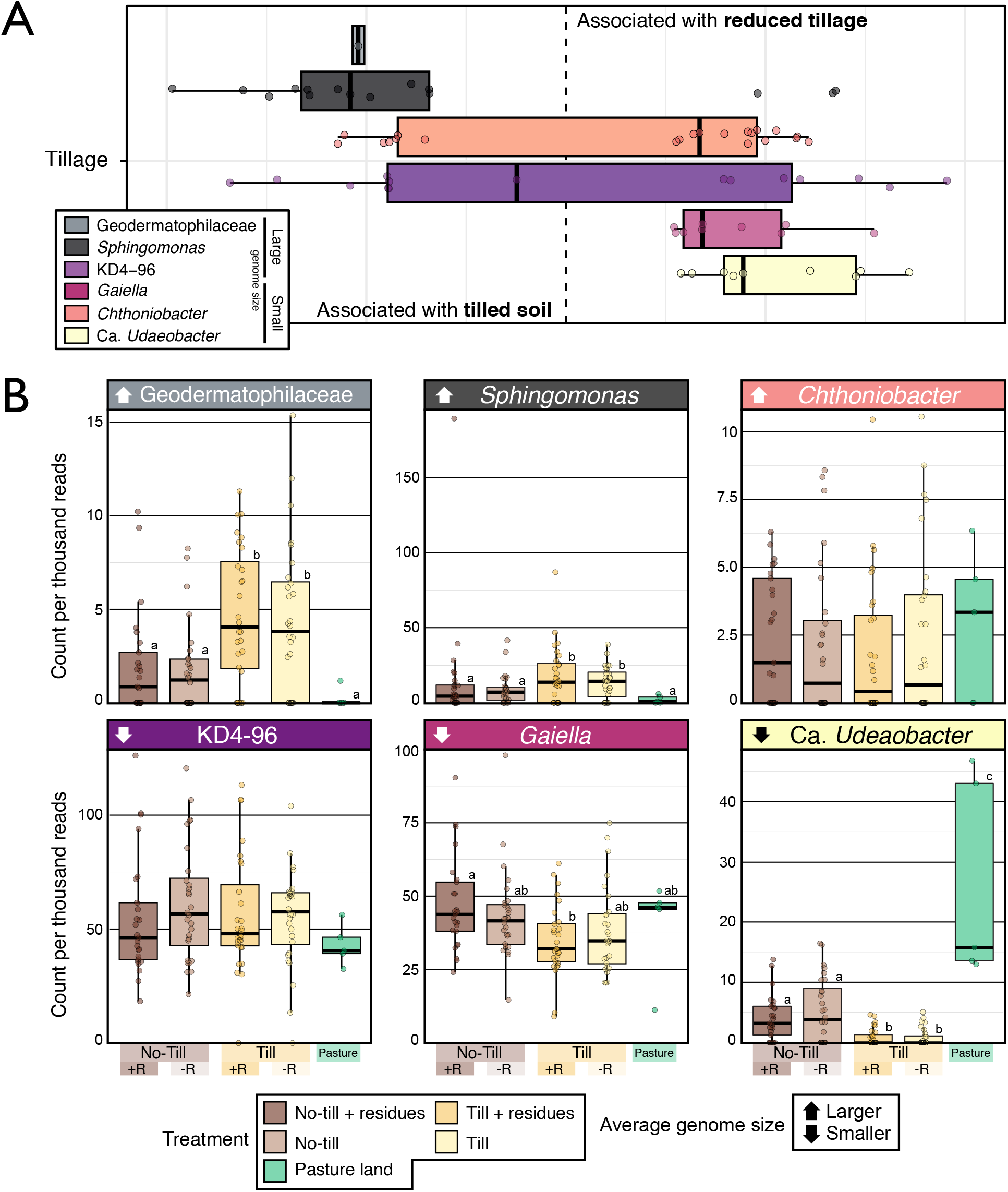
The association of representative taxa from either end of the spectrum of genome size with tillage status were evident in (A) their environment-wide association in the AgroEcoDB and (B) trends in relative abundance in a study of the long-term effects of tillage and plant residue retention, also present in the AgroEcoDB (Koechli, 2016). Taxa with larger genomes were found at higher relative abundances in tilled fields, while the reverse was true for taxa with smaller genomes, consistent with trends in the soil health data. In (B), statistically supported differences are denoted with lettering based on pairwise Tukey HSD tests (p < 0.05).

**Figure S5.**
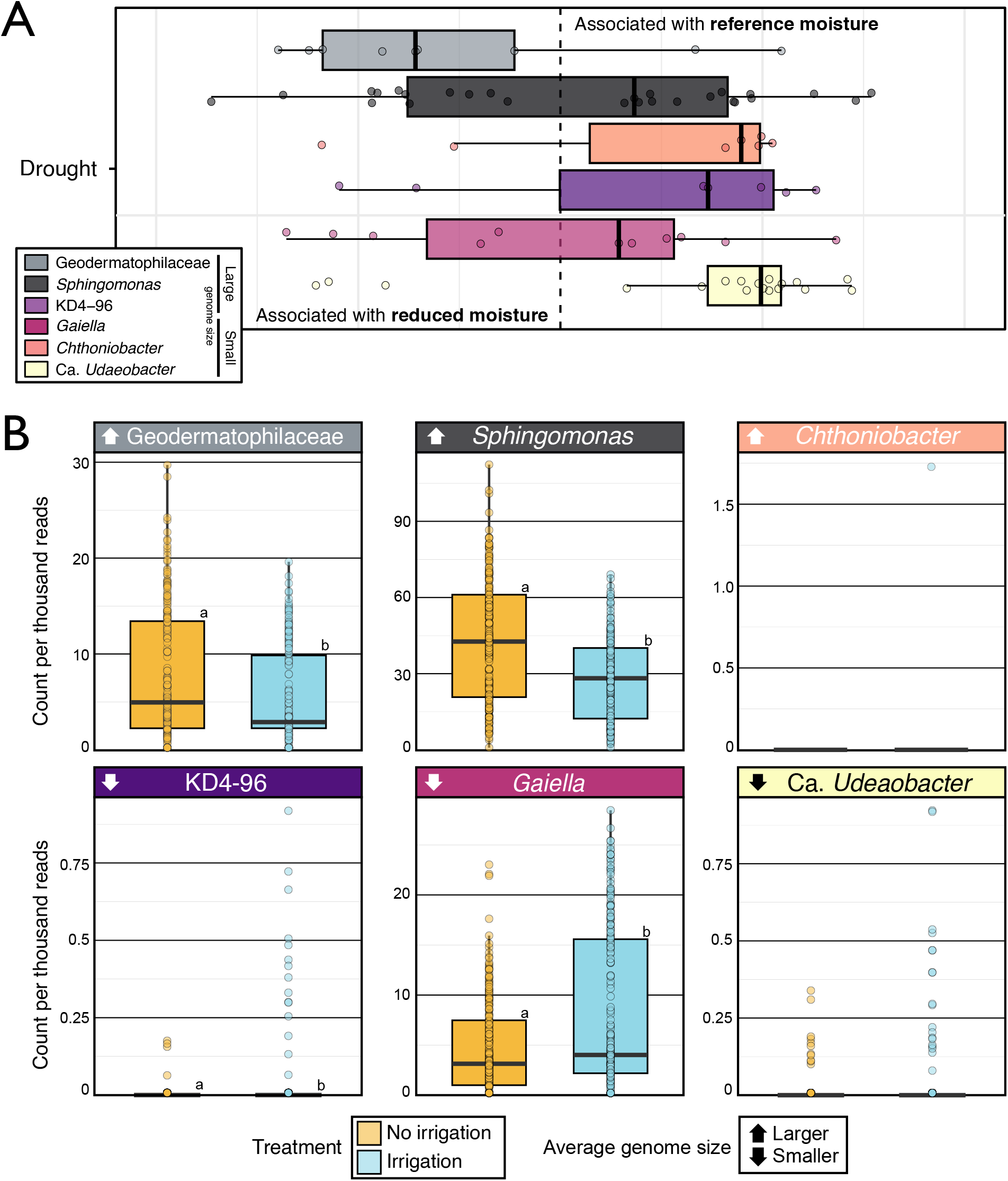
The association of representative taxa from either end of the spectrum of genome size with watering regime was evident in (A) their environment-wide association in the AgroEcoDB and (B) trends in relative abundance in a study of the long-term effects (40 years) of irrigation, also present in the AgroEcoDB (Azarbad et al., 2020). Taxa with smaller genomes were found at higher relative abundance irrigated fields, while the reverse was true for taxa with larger genomes. Statistically supported differences are denoted with lettering based on Wilcoxon tests (p < 0.05).

**Figure S6.**
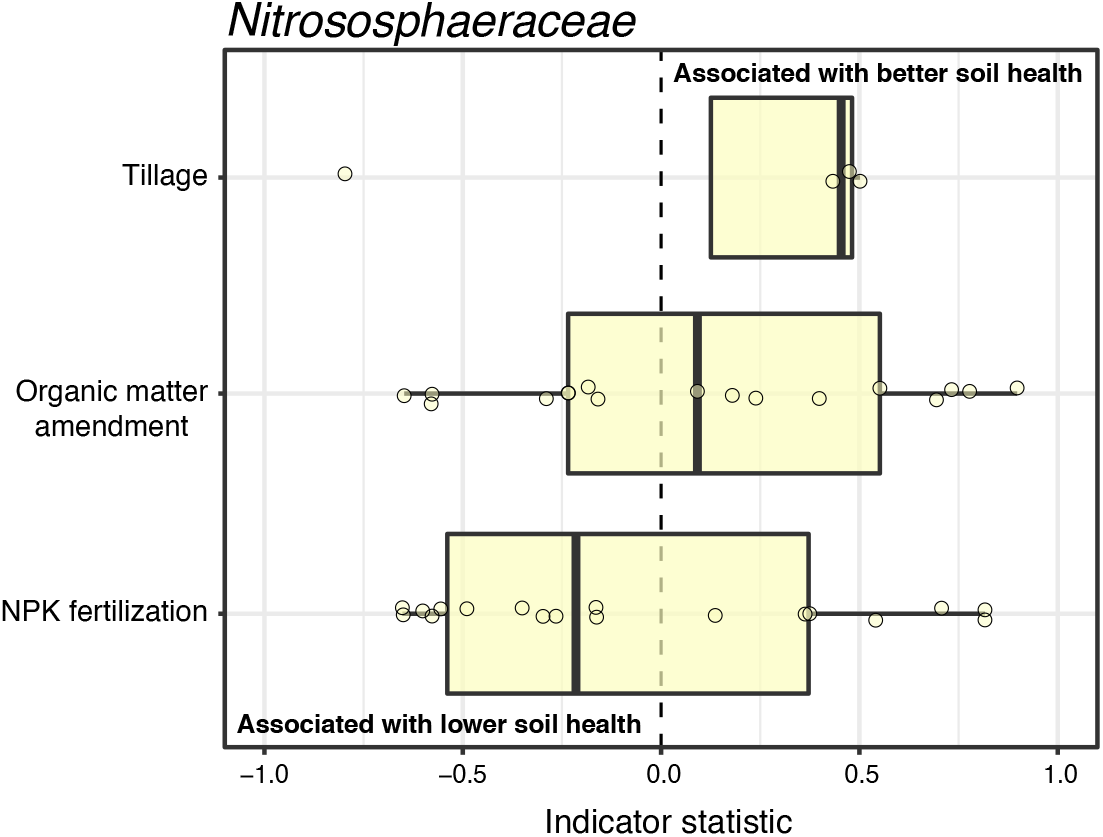
Environment-wide associations of *Nitrososphaeraceae* indicating an association with fertilizer use (‘NPK’).

**Figure S7.**
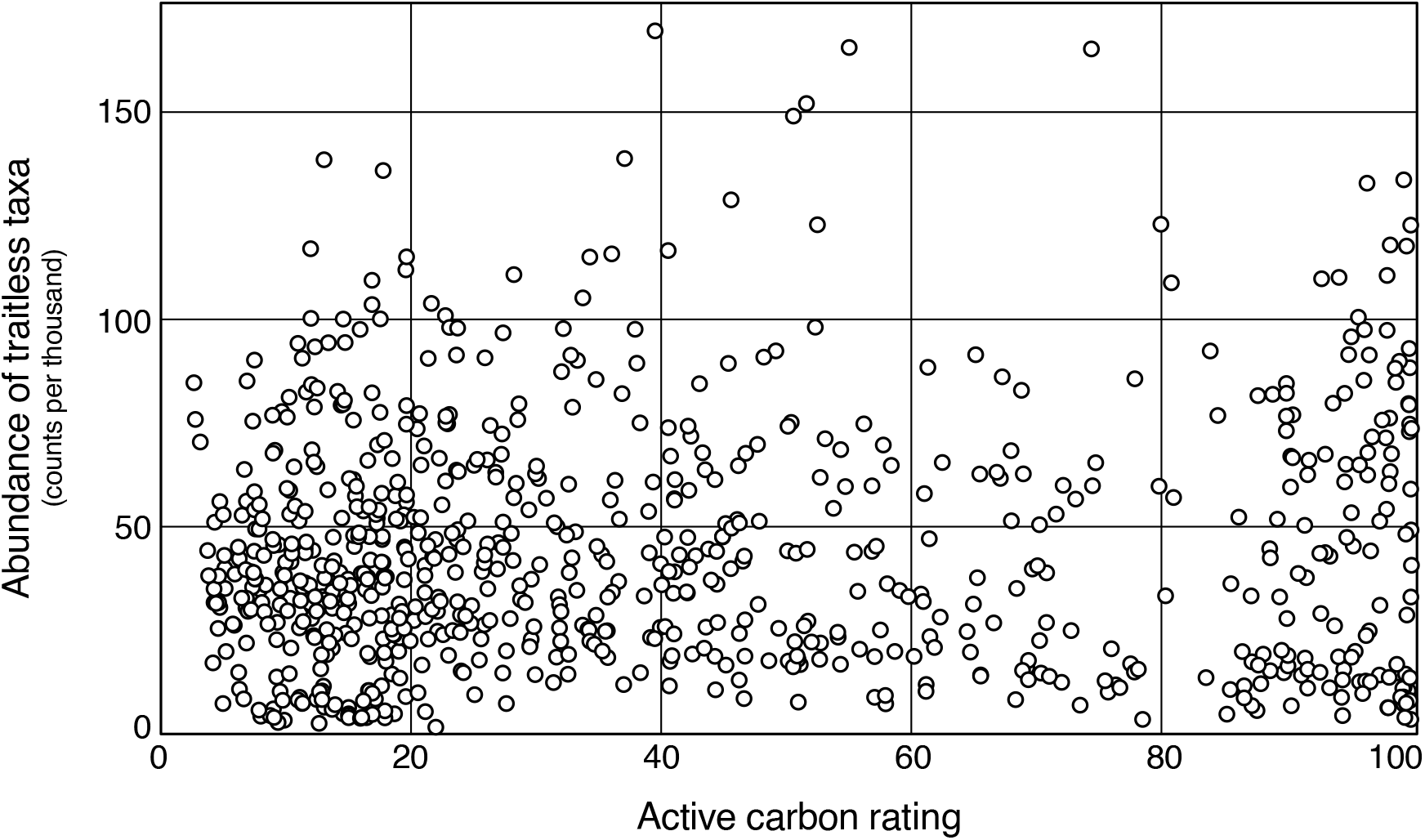
The relative abundance of taxa in amplicon sequencing libraries which could not be assigned a genomic trait. The y-axis corresponds to the aggregate relative abundance of all traitless taxa. There was no correlation with active carbon rating (r = 0.06, p = 0.1).

## Supplementary Information

From Wilhelm *et al*., 2022: “Ecological insights into bacterial indicators of soil health according to their life-history traits and environment-wide associations in agricultural soils.”

### Downloading Sequencing Projects for the AgroEcoDB

All amplicon sequencing projects were downloaded using a series of custom scripts available in the Supplementary Data package, which were executed in the following order: (1) ‘download.SRA.metagenomes’, (2) ‘define.variable.region.py’, (3) ‘download.runs.py’ and (4) ‘get.SRA.metadata.py.’ In brief, these scripts will (1) download sequencing project information for a set of provided NCBI taxonomy IDs, (2) download a subset of runs (n = 10) from each BioProject and determine which variable region of the 16S rRNA gene was targeted using HMMs from VExtractor (Hartmann et al., 2010; included in Supplementary Data) and positional overlap with a reference database, (3) download all sequencing runs from BioProjects which met the criteria (V region etc.) and, finally, (4) download associated study metadata and incorporate into a QIIME2 data object [2]. The identification of overlap with your reference database is critical for ensuring sequences are trimmed to an identical length which is necessary to obtain identical identifiers during ASV calling by DADA2 [3]. In script (3), there is code to ‘walk’ various starting positions to verify the correct information (optional; but recommended). The trimming parameters used for each sequencing project to match soil health data are provided in Table S2. The initial trimming parameters used in preparing soil health data involved removing 5 nt off of both ends of amplicons produced using the standard 515F / 805R primer pair. Script 3 should be run twice: first with option prep_trim_sheet set to ‘Y’. This will output QIIME2 summary ‘qzv’ which should be used to manually verify the sequencing quality of each project using the website: https://view.qiime2.org/. This will also output a ‘trim sheet’ that one can use to input trimming parameters for each project (note: trimming may yield sequence data that no longer overlaps with the reference project). In creating the AgroEcoDB, the number of reads per library was capped at 50,000, since keep all the data from larger sequencing project significantly slowed down the process. All phyloseq objects, representative sequences and indicator species output are available in the Supplementary Data package.

These scripts can recreate analyses will likely be of use to readers wishing to expand on the AgroEcoDB or create their own ASV-based database. However, the scripts are hardcoded in many places and do not serve as an out-of-the-box bioinformatic pipeline. There was considerable effort required to standardize metadata for downstream analyses. For example, identifying and consolidating information about the factor codes used in each experiment / sequencing project. This information is available in the Supplementary Data (‘agroecoDB - study metadata.xlsx’) but be advised that standardizing and cleaning metadata represents the greatest investment in time for building this kind of database. The R script entitled ‘BLOCK00 –Refine metadata’ is provided in the Supplementary Data which provides an idea of the level of manual intervention required.

